# A *CLASP1* variant suggests a phenotypic relation with lissencephaly in humans

**DOI:** 10.1101/2024.03.12.584685

**Authors:** Rawan Alsafeh, Amal AlHashem, Aly Elsayed, Zafer Yüksel, Kalthoum Graies-Tlili, Farah Thabet, Brahim Tabarki

**Author notes:** **Correspondence** to: Dr. Brahim Tabarki, Division of Neurology, Department of Pediatrics, Prince Sultan Military Medical City, P.O. Box 7889, 11159 Riyadh, Saudi Arabia.

## Abstract

Lissencephaly is a severe brain developmental disorder; characterized by reduced brain folding due to defective neuronal migration. This study investigates the genetic basis of lissencephaly in a consanguineous family, focusing on the *CLASP1* gene. Whole-exome sequencing identified a novel homozygous variant (c.4442G>A p.(Arg1481His)) in *CLASP1*. Clinical evaluation revealed severe developmental delays, microcephaly, seizures, and lissencephaly in the affected siblings. The variant was heterozygous in the healthy parents and a heterozygous carrier in an unaffected sibling. This study underscores the role of CLASP1 in brain development and suggests that the identified variant disrupts CLASP1 interaction with the microtubule cytoskeleton, contributing to lissencephaly pathogenesis.

## Background

Lissencephaly is a severe brain developmental disorder; characterized by reduced brain folding due to underlying cortical layering defects. These aberrations arise during embryonic development owing to defective neuronal migration. Progress in molecular genetics has aided the identification of at least 31 lissencephaly-associated genes, with an overall diagnostic yield of over 80% (Koenig, 2021). Many of these genes encode microtubule structural proteins (tubulin) or microtubule-associated proteins, which play major roles in regulating cytoskeleton dynamics during neuronal migration (Romero, 2018). Cytoplasmic linker-associated protein 1 (CLASP1) belongs to a group of proteins known as CLASPs, which are non-motor microtubule-associated proteins that interact with Cap-Gly Domain-containing linker protein (CLIPs), members of the microtubule plus-end tracking protein family (Mimori-Kiyosue, 2005). Considerable evidence now implicates CLASP1 in growth cone orientation, axon guidance, and the regulation of neuronal migration in mice (Sayas, 2019). To date, however, no human monogenic disorder has been associated with the *CLASP1* gene. In this study, we propose a possible phenotypic association between biallelic CLASP1 variants and lissencephaly in humans.

## Materials and Methods

Whole-exome sequencing (WES) analysis was performed in three siblings diagnosed with lissencephaly, along with their parents. We performed this analysis with informed consent from the family, focusing on genes already associated with the clinical manifestations of the patients, including but not limited to lissencephaly, as well as other protein-coding genes not yet associated with a phenotype. The genomic DNAs were fragmented, and the exons of known genes in the human genome, as well as the corresponding exon-intron boundaries, were enriched, amplified, and sequenced simultaneously using Illumina technology (San Diego, CA). The sequenced target regions with an average coverage of 133-fold, achieving a 20-fold coverage for approximately 99% of the regions of interest. We performed short-read whole-genome sequencing, as well as segregation analyses, in this family to rule out other possible genetic etiologies.

## Results

We evaluated three affected siblings from a large consanguineous Saudi family (Figure 1). Table 1 summarizes he main clinical and MRI features. WES revealed a novel homozygous variant, c.4442G>A p.(Arg1481His), in the CLASP1 gene for all three affected siblings (IV-1, IV-2, IV-4) and heterozygous in their healthy, consanguineous parents (III-1, III-2). Sanger sequencing revealed that the unaffected male sibling (IV-3) is a heterozygous carrier for the variant. At birth, all had low weight and microcephaly, with head circumference ranging from 27 to 32 cm (< 3rd percentile), was observed. The patients presented with clinical features of microcephaly (ranging from -3SD to -4SD), early-onset spasticity, intractable epilepsy, and profound developmental delay. All three patients had epilepsy, with seizures starting as early as four months of age. Two patients had generalized tonic-clonic seizures, while the other patient had tonic seizure. Two patients had medically intractable epilepsy, and the condition of one patient was fairly controlled (but he is was relatively younger than the others and with limited follow-up). Neuroimaging revealed multiple brain abnormalities (Figure 2 and 3). MRI in all three patients showed lissencephaly, reduced white matter volume, hypoplastic corpus callosum, and a pontine hypoplasia.

**Table 1.**
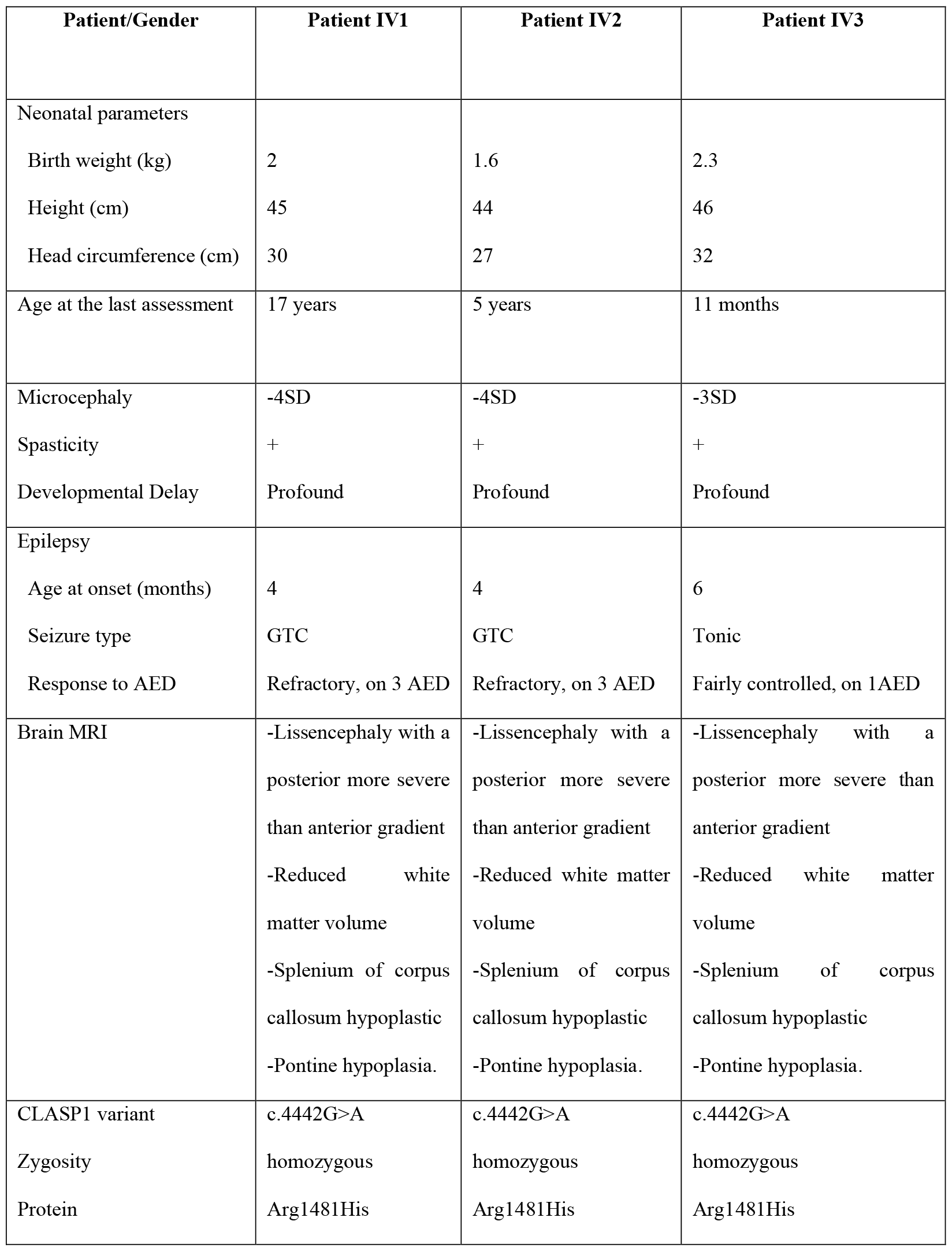
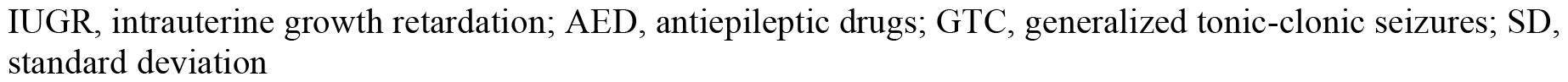
The clinical and genetic characteristic of *CLASP1*-associated neurophenotypes.

| Patient/Gender | Patient IV1 | Patient IV2 | Patient IV3 |
| --- | --- | --- | --- |
| Neonatal parameters |  |  |  |
| Birth weight (kg) | 2 | 1.6 | 2.3 |
| Height (cm) | 45 | 44 | 46 |
| Head circumference (cm) | 30 | 27 | 32 |
| Age at the last assessment | 17 years | 5 years | 11 months |
| Microcephaly | -4SD | -4SD | -3SD |
| Spasticity | + | + | + |
| Developmental Delay | Profound | Profound | Profound |
| Epilepsy |  |  |  |
| Age at onset (months) | 4 | 4 | 6 |
| Seizure type | GTC | GTC | Tonic |
| Response to AED | Refractory, on 3 AED | Refractory, on 3 AED | Fairly controlled, on 1AED |
| Brain MRI | -Lissencephaly with a posterior more severe than anterior gradient<br>-Reduced white matter volume<br>-Splenium of corpus callosum hypoplastic<br>-Pontine hypoplasia. | -Lissencephaly with a posterior more severe than anterior gradient<br>-Reduced white matter volume<br>-Splenium of corpus callosum hypoplastic<br>-Pontine hypoplasia. | -Lissencephaly with a posterior more severe than anterior gradient<br>-Reduced white matter volume<br>-Splenium of corpus callosum hypoplastic<br>-Pontine hypoplasia. |
| CLASP1 variant | c.4442G>A | c.4442G>A | c.4442G>A |
| Zygosity | homozygous | homozygous | homozygous |
| Protein | Arg1481His | Arg1481His | Arg1481His |

IUGR, intrauterine growth retardation; AED, antiepileptic drugs; GTC, generalized tonic-clonic seizures; SD, standard deviation

**Figure 1.**
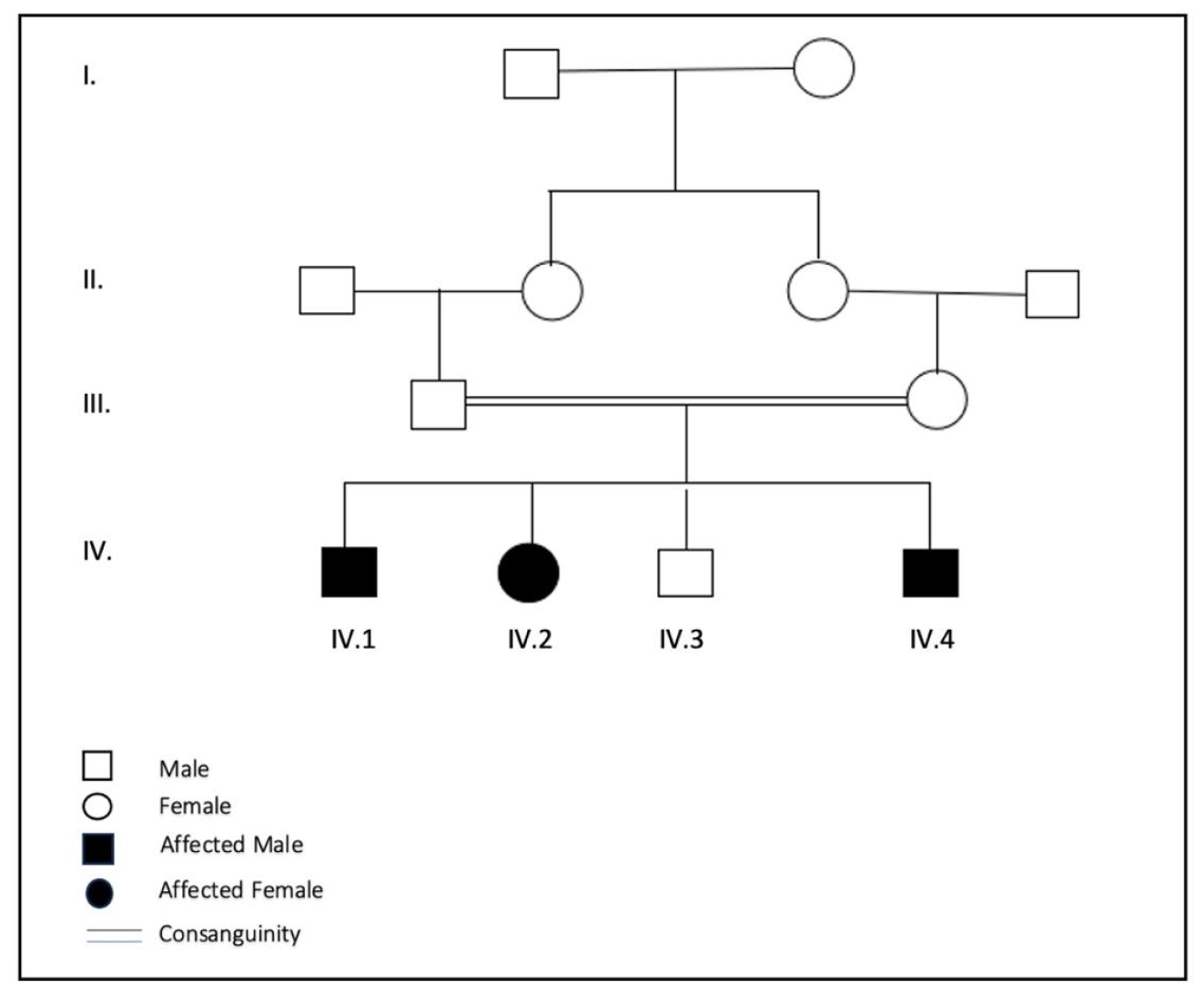
Pedigree of the study family showing the degree of consanguinity between the parents.

## Discussion

Herein, we describe three affected siblings from a large consanguineous family with a novel homozygous variant in the CLASP1 gene. All affected individuals presented early in life with severe to profound developmental delays, primary microcephaly, seizures refractory to antiepileptic treatment, and lissencephaly. Lissencephaly represents a spectrum of malformations in cortical development, including agyria, pachygyria, and subcortical band heterotopia. CLASP1 belongs to a group of proteins known as CLASPs. These proteins are highly expressed in the central nervous system and are non-motor microtubule-associated proteins that interact with CLIPs, a member of the microtubule plus-end tracking protein family that promotes the stabilization of dynamic microtubules in migrating cells (Cushion, 2023). Loss of the CLASP1 C-terminus weakness MT plus-end binding. The clinical and neuroradiological abnormalities observed in CLASP1, such as lissencephaly with a posterior-more-severe-than-anterior gradient, thin/hypoplastic corpus callosum, and pontine hypoplasia, are analogous to the findings seen in *CAMSAP1*-related neuronal migration disorder, phenotype consistent with a severe tubulinopathy (Khalaf-Nazzal). The close alignment of neuroradiological features between the *CAMSAP1*-related neuronal migration disorder and *CLASP1*-related lissencephaly suggests a shared pathomolecular mechanism that underlies these diseases. This observation underscores the significance of the minus end of the microtubule in neuronal migration disorders.

In conclusion, we propose that the biallelic p.(Arg1481His) variant disrupts the interaction of CLASP1 with the microtubule cytoskeleton, leading to lissencephaly.

---

**Figure.**
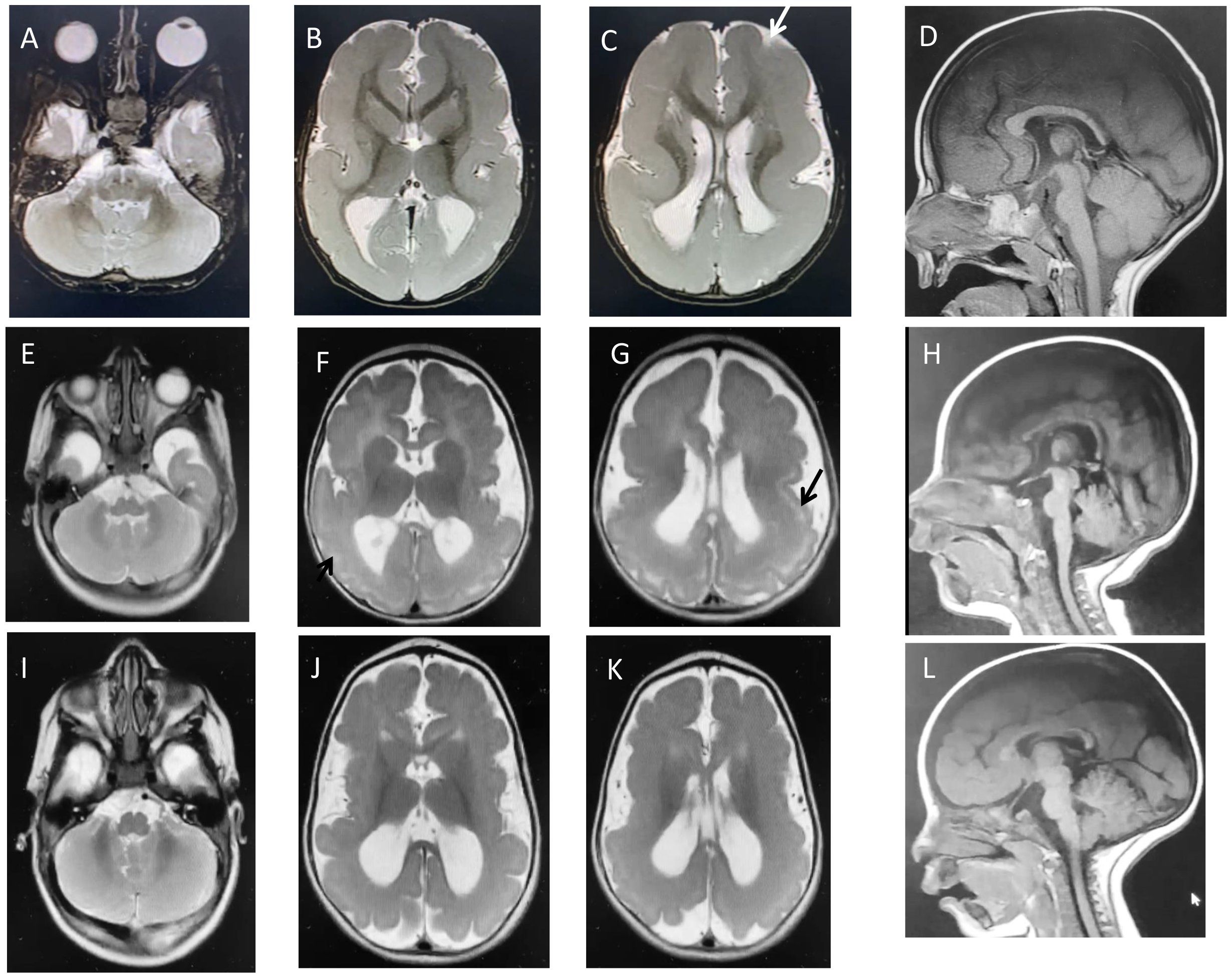
Neuroimaging in three individuals with the *CLASP1*-related lissencephaly: row 1 (A-D) is patient IV:1 aged 4 years. T2-weighted axial images (A-C) show posterior-more-severe-than-anterior gradient with areas exhibiting nearly absent gyration or agyria with extremely shallow frontal sulci (white arrow) and a slightly wide and shallow sylvian fissure. Reduced white matter volume, particularly poorly developed posteriorly, and mild enlargement of the lateral ventricles, mainly posteriorly with a squared pattern. T1-weighted midline sagittal image (D) shows a thin/hypoplastic splenium of the corpus callosum and pontine hypoplasia. Row 2 (E-H) is patient IV:2 aged 2 months and row 3 (I-L) is patient IV:4 aged 9 months. T2-weighted images (E-G and I-K) show diffuse thick cortex with reduced gyration/ pachygyria, slightly more severe posteriorly, along with almost age-appropriate myelination of the reduced white matter. In figures F-G, there is thin T2 hyperintense band within the thick cortex (black arrow) representing a sparse cell layer zone between the thick arrested neuronal layer and the thin superficial molecular layer. Mild enlargement of the lateral ventricles, mainly posteriorly with a squared pattern. T1-weighted midline sagittal images (H, L) show the diffuse hypoplastic corpus callosum and pons.

